# Microfluidic bioprinting of a physiologically relevant thyroid three-dimensional *in vitro* model

**DOI:** 10.64898/2026.04.23.720314

**Authors:** Mirco Sana, Stefan Giselbrecht, Mírian Romitti, Anna M Kip, Sabine Costagliola, Carlos Mota, Lorenzo Moroni

**Affiliations:** Department of Complex Tissue Regeneration, MERLN Institute for Technology-Inspired, Regenerative Medicine, Maastricht University, Maastricht, 6229 ER, The Netherlands; Department of Instructive Biomaterials Engineering, MERLN Institute for Technology-Inspired Regenerative Medicine, Maastricht University, Maastricht, 6229 ER, The Netherlands; Institute of Interdisciplinary Research in Molecular Human Biology (IRIBHM), Université Libre de Bruxelles, 808 route de Lennik, Brussels, 1070, Belgium

**Keywords:** 3D in vitro models, bioprinting, thyroid, endocrine disruptors

## Abstract

Endocrine disruptors (EDs) are an exogenous group of compounds associated with thyroid malfunctioning in the human body. Nonetheless, there are currently no adequate *in vivo* or *in vitro* models for the preclinical testing of these compounds since both animal and two-dimensional (2D) cell-based models are not able to mimic thyroid physiological conditions from both functional and three-dimensional (3D) organization perspective. Recently, bioprinting technologies emerged as an innovative tool in the field of regenerative medicine and advanced 3D *in vitro* models that allow the creation of 3D well-organized structures able to mirror physiologically relevant tissue and organ architectures.

In this study, we evaluated microfluidic bioprinting as a biofabrication technology to develop a 3D *in vitro* model of the thyroid gland. We studied the fundamental parameters to obtain a fine control over the bioprinted fibres for different biomaterials. Then, we assessed the possibility to bioprint single thyroid cells, thyroid spheroids and finally mouse embryonic stem cell-derived thyroid follicles. The different cell types maintained high viability and metabolic activity. The bioprinted thyroid model showed high expression of different early and late functional markers and to be responsive to ED exposure. These bioprinted thyroid constructs could provide a new set of advanced 3D *in vitro* models to test potential EDs and possible adverse outcomes that may be associated with their administration or exposure.

## 1. Introduction

The human thyroid is an endocrine gland, which is normally located in the anterior lower neck. The main functional unit of the thyroid, known as thyroid follicles, are round, hollow structures composed of a single layer of specialized epithelial cells called thyrocytes. The follicles are filled with a fluid known as colloid, which main component is Thyroglobulin (Tg), the precursor of thyroid hormones (THs). The main function of the thyroid is the production of thyroid hormones thyroxine (T4) and triiodothyronine (T3) and their storage inside the follicles. THs are then released in the blood flow and influence a wide range of bodily functions (e.g., control of metabolic rate, energy expenditure, growth, and development).[1–3] A wide range of chemical compounds able to negatively affect thyroid development and functions have been discovered, which are part of the wider group of chemicals capable of interfering with endocrine system known as endocrine disruptors (EDs). In the last decades several international regulators (e.g. the World Health Organization, the European Commission, European Food Safety Authority) have been working in order to identify and regulate these EDs and to investigate their effect on human health and environment [4, 5]. EDs mode of actions (MOAs) are not always known or fully understood, but the most general MOA is related to the EDs ability to bind specific receptors expressed by different tissues causing a downstream effects on their functions [6]. Nonetheless, EDs can interfere over thyroid functionality at different levels, ranging from the impairment of the signaling along the hypothalamic–pituitary–thyroid axis to the direct disruption of TH synthesis. Although many *in vitro* and *ex vivo* assays have been developed to identify EDs and confirm their MOA, many of them suffer several limitations [4]. Classical *in vitro* models are based on 2D cultures of thyroid cells and for this reason they fail to recapitulate the three-dimensional (3D) environment and functionality of the thyroid. Furthermore, 2D culture of cells that are supposed to be in a 3D environment, such as thyroid follicles, could lead to the dedifferentiation of these cells, thus causing a loss of their specific functions.[4] In the last years, different strategies have been studied and developed in order to produce 3D *in vitro* models able to mimic physiological conditions, such as architecture and function. Among these strategies, bioprinting emerged as a promising technology able to produce cell laden 3D biological constructs in a reproducible way. To date, different biofabrication technologies have been developed to offer the best material deposition, each of them with specific advantages and limitations [7, 8].

Here, we used microfluidic bioprinting to exploit its distinctive characteristics for the creation of a hydrogel-based 3D thyroid *in vitro* model. The specific microfluidic bioprinting system used in our study was already assessed in previous works focusing on the bioprinting of soft hydrogels using single cells or organoids representing other tissues [8–10]. Microfluidic bioprinting offers different interesting features for the biofabrication of 3D biological constructs. The bioprinting dispensing system, defined as printhead, is composed of microchannels and valves where bioinks can flow and then be dispensed through a nozzle. Within the channels a coaxial flow is used to focus the crosslinker around the stream of bioink inside the printhead, allowing the crosslink to happen during the printing procedure and the gelled microfibre to be immediately deposited. The pressure applied within the channels, typically between 0 and 500 mbar, is significantly lower compared to conventional extrusion-based bioprinters [11]. The activation of the valves integrated within the printhead allows instead to maintain optimal control over the flow, the switching, and the mixing of the solutions flowing inside the printhead. The possibility to use low viscosity bioinks together with the low pressure applied and the laminar flow within the microfluidic printhead expose the cells to mild printing conditions and minimal shear stress, thus creating a safe bioprinting environment and minimizing the possible causes of cell death [7, 11]. All the characteristics presented above are crucial when addressing the challenges of developing advanced bioprinted models: not only do they allow the production of precise and multi layered 3D biological constructs, but they enable the use of soft materials, which are essential to reproduce models for soft organs. Moreover, bioprinted cells maintained within soft gels will also be able to migrate, proliferate or remodel their microenvironment [12].

In this study, we initially focused on testing the printability of different bioinks and on optimizing the bioprinting settings to produce stable 3D biological constructs with defined geometries. Then, the behaviour of single cells and spheroids inside the different bioinks and the effect of bioprinting was evaluated on their viability and metabolic activity. We then focused on the bioprinting of more complex and physiologically relevant mESCs-derived thyroid follicles. Thyroid follicles’ viability and functional behaviour inside the bioprinted constructs was assessed during the 10 days of culture. Finally, we assessed if the thyroid *in vitro* model was able to respond to the exposure of EDs. Propylthiouracil (PTU) was chosen for its known inhibitive effect on TH synthesis [13] both *in vitro* and *in vivo*. Bioprinted thyroid follicles were exposed to different concentrations of PTU for 10 days and the effect on thyroid functionality was assessed by immunohistochemistry and quantification of TH release.

## 2. Material and methods

### 2.1 Cells expansion and differentiation

*Nthy-ori 3-1 cell culture*. Nthy-ori 3-1 cells, an immortalized human primary thyroid follicular epithelial cell line, was used as they are reported to retain thyroid relevant functions such as iodide-trapping and thyroglobulin production [14–17]. Nthy-ori 3-1 have been used in studies involving growth and control of the human thyroid [14, 18]. Nthy-ori 3-1 cells (Sigma-Aldrich) were cultured following manufacturer guidelines using RPMI1640 with Glutamax + 10% FBS. Nthy-ori 3-1 cells were used for bioink and bioprinting optimization tests. Briefly, Nthy-ori 3-1 cells cultured in monolayer were trypsinized, washed twice with PBS and mixed with the bioink at a final concentration of 4*10^6 cells/mL. When cells were combined with the hydrogels to form the bioinks, medium was supplemented with Penicillin/Streptomycin 1:1000. *Nthy-ori 3-1 spheroid production.* Nthy-ori 3-1 cells spheroids were generated using thermoformed microwell arrays as previously described [19, 20]. Briefly, arrays containing microwells with a diameter of 250 µm were produced form polycarbonate films. The formed membranes were washed and sterilized by immersing them in 70% Isopropanol under agitation. Microthermoformed membranes and O-rings (ERIKS, EPMD) were washed in consecutive isopropanol solutions of 70%, 20%, and 2x 0% for 10 min to completely wet the microstructured surface. Membranes were placed in 24-well plates, secured with O-rings, and residual air bubbles were removed through washing with PBS. Each well was then covered with 250 µL of Pluronic F-108 3% (w/v) (Sigma-Aldrich) in MilliQ water, previously sterile filtered. Membranes were incubated overnight. The day after, membranes were washed with PBS and a nThy ori 3.1 cell suspension was seeded in order to aggregate the cells in the microwells to spheroids overnight. The day of bioprinting, spheroids were observed under the microscope and pictures were analyzed using ImageJ 1.52p software to assess their diameter. Two different concentrations of spheroids were tested for bioprinting: 2000 and 4000 spheroid/mL.

*mESC-derived thyroid organoid*. Recombinant murine ESC line (A2Lox Nkx2-1-Pax8), generated as previously described [21, 22], were firstly aggregated into embryonic bodies (EBs) and subsequently differentiated into thyroid follicles. The schematic of the differentiation protocol is shown in **Figure 3A**. Briefly, mESCs were cultured on γ-ray irradiated mouse embryonic fibroblast (MEF) feeders in DMEM supplemented with 15% ES Cell qualified FBS (Sigma Aldrich, St. Louis, USA), IK0701 LIF (1000 U mL−1) (ORF Genetics, Kopavogur, Iceland), nonessential amino acids (0.1 × 10−3 m final), sodium pyruvate (1 × 10−3 m), penicillin and streptomycin (50 U mL−1 final), and 2-mercaptoethanol (0.1 × 10−3 m). mESCs were then collected and maintained as hanging drops (1000 cell/drop) in order to produce EBs using differentiation medium containing DMEM supplemented with 15% FBS, vitamin C (50 µg mL−1), nonessential amino acids (0.1 × 10−3 m), sodium pyruvate (1 × 10−3 m), penicillin and streptomycin (50 U mL−1), and 2-mercaptoethanol (0.1 × 10−3 m). EBs were collected after 4 days and mixed with Matrigel Growth Factor Reduced (354230, Corning, New York, USA). 50 uL Matrigel drops were then plated into 12-well plates, each droplet containing around 30 EBs. Matrigel droplets were cultured for 3 days using differentiation medium supplemented with Doxycycline (1 µg mL−1), followed by 14 days in differentiation medium supplemented with 8-Br-cAMP (10 × 10−6 m, B 007, Biolog, Hayward, USA). Cell differentiation was monitored during culture by observing cell morphology and bovine Tg promoter (bTg)-driven GFP expression.

### 2.2 Hydrogel screening

An initial hydrogel screening was performed envisioning the bioink formulations for bioprinting. AG10 Matrix ™ (AspectBiosystems), alginic acid sodium salt from brown algae (low viscosity, Sigma-Aldrich), gelatin solution Type B, tissue culture grade (Sigma-Aldrich), gelMA 300 Bloom with 60% degree of substitution (DS) (Sigma-Aldrich), lithium phenyl-2,4,6-trimethylbenzoylphosphinate (LAP) photoinitiator (Sigma-Aldrich), fibrinogen from bovine plasma Type I-S, 65-85% protein (Sigma-Aldrich), were used as hydrogel precursors either individually or in combined formulations.

All bioinks have been produced by dissolving them in PBS and then sterile filtered using a 0.2 µm syringe filter. The crosslinker solution used in our experiments had this final composition: 125 mM calcium chloride (anhydrous, granular, ≤ 7.0 mm, ≥ 93.0%; Sigma) + 2% poly(vinyl alcohol) (PVA, average Mw 85,000-124,000, 87-89% hydrolysed; Sigma) in MilliQ water. When fibrinogen containing bioinks were used, the crosslinker solution was supplemented with thrombin active (High Activity) from bovine plasma (Sigma-Aldrich) for a final concentration of 5 U/mL. When GelMA based bioinks were used, the produced hydrogels were irradiated with UV light at 450nm with an intensity of 10 mW/cm2 for 60 seconds by using a LEDD1B -T-Cube LED Driver (Thorlabs). The final composition of all bioinks used are listed in Table 1, together with the codes used to refer to each bioink composition within this work. The properties of these materials are described in Supplementary Table S1.

**Table 1.** Bioink formulations tested for microfluidic bioprinting and bioink identification code used.

| **Bioink composition** | **Bioink code** |
| --- | --- |
| AG10 Matrix™ | **AG10** |
| Alginate 4% w/w | **AG4** |
| Alginate 2% w/w | **AG2** |
| Alginate 1, 5% w/w + Gelatin 0, 5% v/w | **AGg** |
| Alginate 2%w/w + Fibrinogen 1mg/ml | **AGf1** |
| Alginate 2%w/w + Fibrinogen 5 mg/ml | **AGf5** |
| Alginate 2% w/w + GelMA 1% v/w | **AGge1** |
| Alginate 2% w/w + GelMA 2% v/w | **AGge2** |

### 2.3 Bioink droplets production

In order to test both metabolic activity and cell viability in different bioink formulation, nThy-ori 3-1 cells were mixed with different bioinks (AG10, AG4, AG2, AGg, AGf1, AGf5, AGge1, AGg2) with a final concentration of 4×10^6^ cells/mL. In order to produce droplets of bioink containing encapsulated nThy-ori 3-1 cells, 20 µL of each bioink (total number of cells inside the droplet: 8×10^4^) was deposited in an untreated well plate and covered with 125 mM calcium chloride crosslinker solution for 10 minutes. The crosslinker solution was supplemented with thrombin when fibrinogen containing gels were used, while GelMA containing hydrogels were also crosslinked by UV light for 1 minute using a wavelength of 405 nm and an intensity of 2,50 mW/cm^2^. Then, the crosslinker solution was removed and the droplets were cultured in RPMI 1640 with GlutaMAX (Thermo Fisher Scientific) supplemented with 10% FBS and P/S (100 U/ml). The cell medium was refreshed every two days. The crosslinking procedure was repeated on day 3 of culture to maintain the gel stable over time. At day 1, 3 and 7 of culture, droplets were randomly collected to assess viability and metabolic activity.

### 2.4 Microfluidic bioprinting

A microfluidic bioprinter (RX1™, Aspect Biosystems, Canada) equipped with a DUO-1 Printhead () was used. The bioprinter presents 4 pressurized reservoirs (two containing the different bioinks, one containing a crosslinker solution and one containing a buffer solution) which were connected by a tubing system to the printhead. The printhead comprises of pressure-actuated valves that can be opened or closed in order to allow biomaterials, crosslinker and buffer solutions to flow within the printhead. The flow and the ratio between each solution was controlled by applying different pressures to each reservoir. The bioink was extruded simultaneously with the crosslinker coaxially to ensure a uniform crosslinking inside the printhead and to prevent the direct contact of the bioink with the walls of the nozzle. The obtained bioink filament was then extruded from the nozzle of the printhead onto a vacuumed insert that removed the excess crosslinker solution and any uncrosslinked material.

The RX1 printer was placed inside a Biosafety Cabinet to ensure sterility of the bioprinted constructs. All the materials listed above were autoclaved before using in order to assure sterility and avoid risks of contamination

### 2.5 Bioprinting parameter assessment

To analyze the ability of the microfluidic bioprinter to tune the diameter of the produced fibres, different pressure settings have been tested on different bioink formulations. Our goal was to produce fibres with a diameter of around 200 µm and to be able to deposit the fibres with a good precision. The procedure in order to print with this specific bioprinter was extensively described elsewhere [8].The pressure applied is indicated as 20-10-100-90 mbar. The first value refers to the pressure applied to material 1 (the bioink used to produce the fibre), the second number refers to the pressure applied to material 2 (not used), the third value refers to the pressure applied to the crosslinker solution, and the fourth value refers to the pressure applied to the buffer solution. The fibres produced with each setting were collected and transferred into a petri dish. For each condition analyzed, a minimum of three fibres were produced from the same bioink batch. For each fibre, several images were taken to cover all its length. Each picture was analyzed with ImageJ software. A minimum of three measurements were acquired for each photo to obtain the average dimension of the fibre and the standard deviation.

### 2.6 3D bioprinted constructs production

To assess cell behavior after bioprinting with a RX1™ Bioprinter, we optimized the production of a standardized bioprinted construct and assessed its stability over time. By using the *Aspect Studio* software integrated in the bioprinter, an 8 x 8 x 2 mm rectangular construct was designed with an infill of 30% and two perimeters at the outer border of the structure and adjacent to each other. The two perimeters were added to increase stability of the structure. Bioprinting speed was adjusted between 30-35 mm/s to obtain the best fibre deposition. The pressures applied during the bioprinting procedure were tuned to obtain a fibre of 200 µm. Bioprinted constructs were printed on a Thincert cell culture insert for 12 well plates, pore diameter 8µm (Greiner), and cultured on the insert. Constructs produced were covered with crosslinker solution for 15 minutes after printing and then culture medium was added. The cell medium was changed every two days. The crosslinking procedure was repeated also on day 3 of culture to maintain the constructs stable over time. At day 1, 3 and 7 of culture, bioprinted constructs were collected to assess viability and metabolic activity as described below.

### 2.7 Viability assay

Cell viability inside hydrogel droplets and bioprinted constructs was assessed at different time points using a LIVE/DEAD viability/cytotoxicity Kit (Thermofisher) based on Calcein-AM, which stains live cells green, and ethidium homodimer-I (EthD-I), which stains dead cells red. Quantification of live/dead fluorescence signal was performed using ImageJ software, which allows to get an estimate of the area occupied by alive cells (Green area) and the area occupied by dead cells (Red area). An evaluation of the percentage of alive cells was obtained using the formula: % Viability= Green area/(Green area + Red area).

### 2.8 Metabolic activity assay

To assess cell metabolic activity inside hydrogels two different methods described below were used. *PrestoBlue™ Cell Viability Reagent (Thermo Fisher Scientific)*. PrestoBlue assay was used to evaluate mitochondria respiration through the reduction of a resazurin-based solution by cells into resorufin; metabolic activity was evaluated quantitatively through fluorescence measurements (Excitation⁄Emission wavelengths: 560nm⁄590nm). For 3D hydrogels samples (droplets and bioprinted constructs), culture media was removed and samples washed twice with PBS. Each sample was covered with PrestoBlue™ solution diluted in culture medium (1:10). Samples were incubated at 37°C for 2 hours covered in aluminum foil to protect them from light. After incubation, 100uL of Presto Blue solution were collected, transferred into a black 96 well plate and analysed with CLARIOstar® Plus plate reader.

*CellTiter-Glo® 3D Cell Viability Assay (Promega)*. This assay is based on a thermostable recombinant luciferase that is used to assess the ATP level specifically in 3D cell cultures by analyzing the luminescent signal produced by the reaction between luciferase and ATP. The assay was performed following the manufacturer’s instruction. Hydrogels were washed with PBS and transferred in a 96 well plate. Each sample was covered with 100 μL of CellTiter-Glo® 3D Reagent and 100 μL of cell culture media. The well plate was then placed on a plate shaker at the lowest speed for 5 minutes to assure the penetration of the working solution inside the droplet, induce cell lysis and improve ATP release form the 3D sample. The plate was then incubated for 25 minutes at room temperature protected from the light. Finally, the luminescence level was recorded using CLARIOstar® Plus plate reader following manufacturer instructions.

### 2.9 mESCs-derived thyroid follicles bioprinting and ED exposure

mESCs-derived follicles were obtained as described above. The day of bioprinting mESCs were recovered from Matrigel prior mixing with bioink AGf5. Briefly, Matrigel droplets were digested using a solution of Dispase II (10 mg/ml in HBSS) and Collagenase type IA (1.37 mg/ml in HBSS), and follicles were extracted and resuspended in media. Follicles were selected based on their dimensions by two consecutive filter passages using a 100 µM and a 30 µM cell strainer. Follicles were then mixed with the bioink with a final concentration between 35-40.000 follicles/mL. A woodpile structure was bioprinted using the software of the bioprinter with these parameters: 1.5 x 8 x 8 mm dimensions, infill 30%, speed 35 mm/s, layer width 0.22 mm, layer height 0.11 mm, for 13 layers in total. The constructs had also a 2 layers perimeter. Bioprinting parameters were assessed to increase printing precision and the filament produced was expected to be around 200 µm. Bioprinted samples were randomly divided into three groups and cultured with media supplemented with Propylthiouracil (PTU, Sigma Aldrich). PTU is a known ED able to prevent thyroxine (T4) and triiodothyronine (T3) hormone synthesis by inhibiting the enzyme thyroid peroxidase, which converts iodide to an iodine molecule and incorporates the iodine molecule into amino acid tyrosine [23]. To prepare PTU containing media, PTU powder was dissolved in pure DMSO to prepare a stock solution of 400 mM PTU (100% DMSO). PTU stock solution was diluted using differentiation media to obtain three different concentrations: CONTROL (DMSO), HIGH PTU (2 mM) and LOW PTU (10 µM). The three conditions were cultured in parallel for a total of 10 days. Media was refreshed every two days after bioprinting. A post-crosslinking procedure was repeated also on day 3 of culture with a 125 mM CaCl2 solution for 10 minutes to increase constructs stability. On Day 1, 7 and 10 after bioprinting, constructs were analyzed to observe follicles viability and distribution. For each condition, one construct was washed with HBSS and incubated with EthD-1(1:1000) and Hoechst Solution (1:1000) in HBSS for 30 minutes. Constructs were then immediately analyzed at the microscope.

### 2.10 Immunostaining characterization

Hydrogel droplets and bioprinted constructs have been collected at different time points (day 1, 7, and 10), washed twice with HBSS and fixed with PFA 4% for 1 hour and room temperature. After fixation, samples were washed again with HBSS and stored at 4°C until needed. On the day of the immunostaining, samples were washed twice with HBSS on an orbital shaker at the lowest setting for 10 minutes. Samples were then covered with a Blocking/Permeabilization solution with a final composition of 3% w/v BSA, 5%v/v goat serum, 0,2% TritonX prepared in HBSS. Samples were incubated for 1 hour at room temperature on a shaking plate Primary antibodies were diluted in HBSS 3% BSA, 1% Goat serum/donkey serum, 0.1% TritonX solution. Blocking/Permeabilization solution was completely removed from the samples and each sample incubated in primary antibody solution and 4 C° overnight. The day after, secondary antibodies were diluted in the same solution used for primary Ab. The primary antibody solution was removed, samples were then washed three times with HBSS for 10 minutes on the orbital shaker and incubated for 2 hours at room temperature with secondary antibodies solution, protected from light. After washing three times for 15 minutes with HBSS, samples were incubated with DAPI solution (1:100 in HBSS) for 15 minutes at room temperature and a final washing with HBSS was performed for 10 minutes. Samples were then transferred on a glass-bottom petri dish to be analysed by optical microscopy (Light Microscope Leica TCS SP8 STED). The following antibodies and dyes were used: Alexa Fluor™ 488 Phalloidin (Thermo-Fisher, 1:100) 4′,6-Diamidino-2-phenylindole dihydrochloride (DAPI) (Sigma-Aldrich,1:100), rabbit anti-Thyroglobulin (A0251 Dako, 1:300 and 1:1000) mouse anti-ZO-1 Monoclonal Antibody (ZO1-1A12) (Invitrogen, 1:300), mouse anti-L-Thyroxine T4 (Thermo-Fisher 1:100).

### 2.11 Evaluation of Thyroid hormone production

Culture media from every culture condition (Control, PTU 2 mM, PTU 10 µM) was collected at different time points and stored at -80C° until further analysis. At the end of the culture time, the collected media samples were used to quantify the amount of T4 released in the culture media over 10 days of culture from the bioprinted constructs. To do so, we used a Thyroxine (T4) Competitive ELISA Kit (ThermoFisher scientific), for each condition tested with the media collected from at least three different samples in duplicates and from two different experiments. T4 concentration was normalized over cell free media and fold change was measured over the average of the Control (CTRL) group.

### 2.12 Statistical analysis

Statistical significance was calculated using Student’s t-test. A P value smaller than 0.05 (P < 0.05) was considered statistically significant (∗). For T4 quantification analysis, two different sets of bioprinted follicles were used for a total of seven samples for each condition tested.

## 3. Results

### 2.1. Hydrogels screening: cytocompatibility and bioprintability

*Nthy-ori 3-1 viability and metabolic activity*. To select the best bioink composition to produce a bioprinted thyroid model, Nthy-ori 3-1 were mixed with several alginate based bioinks and their behavior inside hydrogel droplets was assessed by observing cell viability, morphology and metabolic activity. When Nthy-ori 3-1 were cultured inside hydrogel droplets, cell viability remained stable around 70% for all compositions tested (AG10, AG2, AGg, AGf1, AGf5, AGge1, AGge2) during the first week of culture (**Supplementary Figure S1A**). When observing metabolic activity using Presto Blue assay, mitochondrial activity decreased after 7 days for every hydrogel composition used (AG4, AG2, AGg, AGf1, AGf5, AGge1, AGge2) **(Supplementary Figure S1B)**. On the contrary, when Cell Titer-Glo® 3D Cell Viability Assay was used, ATP level of the encapsulated cells remained stable over time during the first week of culture **(Supplementary Figure S1C).** It is important to highlight that, in all conditions tested, cells were initially deposited as single cells inside hydrogels, but they were able to autonomously form clusters over the first week of culture, thus proving their ability to remodel their 3D environment after bioprinting. (**Supplementary Figure S1A and Supplementary Figure S4A)**. While these results did not provide an overall best candidate bioink for our model, we decided to focus our screening on two specific bioinks, namely AGg and AGf5 to exploit the biological features of gelatin and fibrinogen together with the robustness offered by alginate. *Microfluidic bioprinting evaluation.* In parallel to the cytocompatibility tests, bioprintability of the different bioinks was also tested with the goal of producing both fibres and 3D constructs. The main objective was to produce fibres with a diameter of around 200 µm, thus below the known diffusion threshold to guarantee diffusion of oxygen and nutrients if vasculature is absent and prevent cell apoptosis [24–26]. Both AGg and AGf5 could be bioprinted and the diameter of bioprinted fibres could be tuned by modifying the pressure applied inside the printhead (**Figure 1B**). By increasing the pressure applied to the material channel, the amount of bioink flowing inside the printhead increased, thus increasing the produced fibre diameter. By changing the applied pressure from 20 to 60 mbar, the obtained fibre diameter using AGf5 increased from 118 µm ± 21 µm to 218 µm ± 22 µm. When testing AGg, an increase of pressure from 10 mbar to 50 mbar caused an increase in diameter from 217 µm ± 17 µm to 321 µm ± 24 µm. When the pressure applied to the crosslinker channel was modified, an increase in pressure caused a reduction of the fibre diameter. When bringing the crosslinker pressure from 80 mbar to 120 mbar, the diameter of fibres produced using AGf5 decreased from 200 µm ± 29 µm to 139 µm ± 25 µm while the diameter of fibres produced using AGg bioink reduced from 262 µm ± 24 µm to 222 µm ± 27 µm. Similar results were obtained with all other tested bioink compositions (**Supplementary Figure S2A**). The effect of bioprinting speed on fibre diameters was also investigated **(Figure 1B)**: AGf5 fibre diameter varied from 152 µm ± 20 µm to 149 µm ± 26 µm when bioprinting speed was increased from 5 mm/s to 25 mm/s. AGg fibre diameter varied from 200 µm ± 32 µm to 179 µm ± 23 µm when bioprinting speed was increased from 20 mm/s to 30 mm/s. In both cases, differences were not significant. Speed variation was not found to have an influence on fibre dimension in all tested bioink (**Supplementary Figure S2C),** although higher printing speeds were correlated with a better fibre deposition and necessary to avoid the fibre coiling phenomena that could be observed when fibre extrusion happened faster compared to fibre deposition (**Supplementary Figure S2B).**

**Figure 1.**
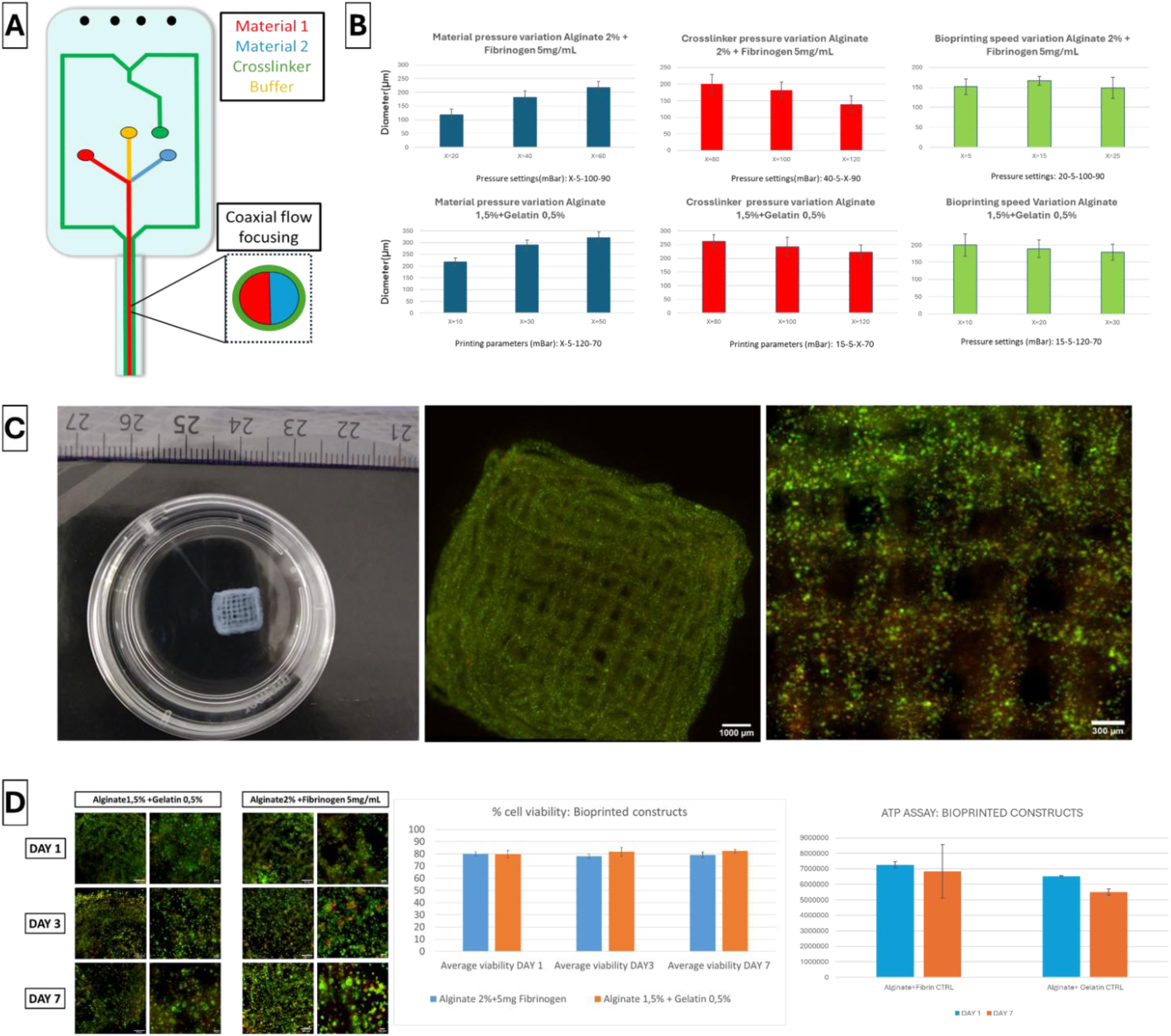
Bioprinting parametric optimization and single cell bioprinting. **(A)** Schematic of microfluidic DUO-1 printhead. **(B)** Fibre average diameter obtained by tuning the pressure applied to the material (BLUE) or to the crosslinker (RED) solution and bioprinting speed (GREEN) using AGf5 and AGg bioinks. **(C)** Macroscopic view of a bioprinted woodpile structure with a designed dimension of 10×10×3mm (width/length/height)- n. layers:30, infill 30%, bioprinting speed 35 mm/s produced using AGf5 bioink and cell viability within the fibres (Scale Bar: 1000 µm and 300 µm). **(D**) Cell viability assessed using LIVE/DEAD viability/cytotoxicity Kit and ATP levels assessed using CellTiter-Glo® 3D Assay of Nthy-ori 3-1 from day 1 to day 7 inside bioprinted constructs produced using AGg and AGf5 bioinks (Scale Bar: 500 µm and 100µm).

In conclusion, it was possible to produce fibres with diameters ranging between 150 and 300 μm with different alginate-based bioinks. Most importantly, pressures applied inside the printhead during the bioprinting procedure ranged between 20 and 150 mbar, are considerably lower compared to other classic bioprinting techniques such as extrusion or inkjet bioprinting [24, 27]. These mild bioprinting conditions should prevent cell death during the bioprinting procedure and together with the tunable fibre diameter and the subsequent exchange of oxygen and nutrients should provide a favourable environment for bioprinted cells.

### 2.2. Bioprinted thyroid constructs

*Nthy-ori 3-1 single cell bioprinting.* The optimized printing parameters were used to produce a more complex bioprinted structure for the creation of the 3D *in vitro* model. A woodpile structure was produced using AGf5 and AGg bioinks mixed with Nthy-ori 3-1 cells. It must be underlined that since bioprinting is performed in wet conditions using low viscosity bioinks, construct geometry might be progressively altered while producing bigger structures. Bioprinted constructs remained stable up to two weeks after bioprinting and an optimal cell distribution inside the bioprinted constructs was achieved (**Figure 1C**). It was possible to produce several constructs (up to 30) per bioprinting session, thus proving throughput efficiency of the microfluidic bioprinting technique. Cell viability remained stable during the first week of culture after bioprinting and, most importantly, the percentage of live cells was assessed around 80% for both bioprinted bioinks (**Figure 1D**) in comparison to the 70% cell viability observed in hydrogel droplets. These results could be due to the improved geometry of the bioprinted constructs, which allows a more even distribution of the cells and a better diffusion of oxygen and nutrients inside bioprinted fibres compared to the droplets. When assessing ATP levels in bioprinted constructs, a decrease in ATP content was observed during the first week of culture when the AGg bioink was used, but not when the AGf5 bioink was used. It is important to highlight how also inside bioprinted constructs cells were initially dispersed as single cells inside hydrogels, but they were able to autonomously form clusters over the first week of culture (**Figure 1D**).

These results show that, after bioprinting, cells remained not only alive and metabolically active, but they were also able to remodel the hydrogel in order to aggregate to clusters. These findings altogether underline that the chosen microfluidic bioprinting technique and the developed alginate-based bioinks allow the production of bioprinted structures with precise geometries and good resolution while using low viscosity materials with a fast-crosslinking procedure that does not influence negatively cell behavior. At the same time, our alginate-based bioinks AGg and AGf5 provided encapsulated cells with an optimal microenvironment able to offer at the same time mechanical support and to promote the creation of more complex structures post-bioprint. In light of these findings, we decided to rely on AGf5 bioink for the following tests for the creation of a thyroid *in vitro* thyroid model.

*Nthy-ori 3-1 spheroids bioprinting.* To emulate the physiological arrangement of follicles in a thyroid, the possibility of bioprinting spheroids obtained from Nthy-ori 3-1 cells was investigated. Spheroids were produced using microthermoformed membranes. The spheroid average diameter was assessed at 173 µm ranging from a minimum of 117 µm to a maximum of 254 µm. It was nonetheless possible to bioprint spheroids of every dimension mixed with AGf5 bioink at two different concentrations (2000 follicle/mL and 4000 follicles/mL). Spheroids could be extruded without problems; no clogging was observed within the microfluidic printhead and spheroid distribution was homogenous within the bioprinted constructs (**Figure 2A**). Most importantly, spheroid morphology was not affected by the bioprinting procedure and spheroids maintained a high viability inside the bioprinted constructs for a week after bioprinting for both concentrations tested (**Figure 2B** and **Supplementary Figure S3)**. In particular, the viability assay did not show the presence of a necrotic core inside the bioprinted spheroids. In conclusion, microfluidic bioprinting further proved its ability to bioprint more complex structures compared to single cells, without presenting technical issues and without negatively affecting the bioprinted spheroids.

**Figure 2.**
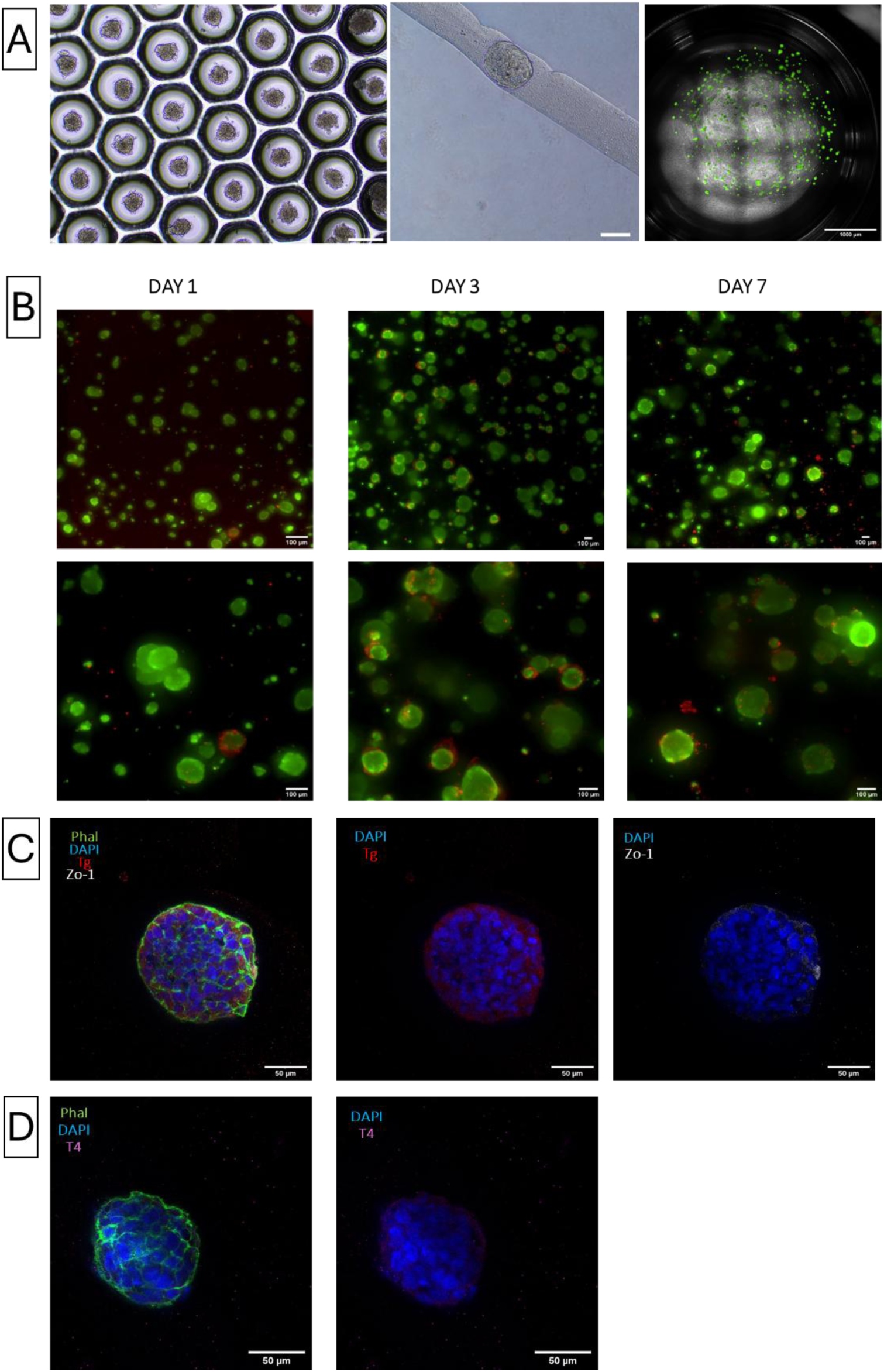
N-thy-ori-3.1 spheroid bioprinting. **(A)** N-thy-ori-3.1 spheroids produced using thermoformed microwell array (Scale bar:300 µm) and a spheroid extruded inside a bioprinted fibre produced using microfluidic bioprinting (Scale Bar:100µm). Macroscopic view of spheroid distribution inside bioprinted constructs produced using AGf5 with a final spheroid concentration of 4000 spheroids/mL (Scale bar: 1000 µm). **(B)** Nthy-ori 3-1 spheroids viability assessed at day 1, 3 and 7 after bioprinting using a LIVE/DEAD viability/cytotoxicity assay. Live cells-green, dead cells-red (Scale bar: 100 µm). Spheroid immunostaining with different thyroid markers at day 7 after bioprinting: (C) Thyroglobulin (Tg)-red; Tight junction protein 1 (ZO-1)-white;(D) Thyroxine (T4)-pink; Phalloidin-green; DAPI-blue (Scale Bar:50µm)

*Nthy-ori 3-1 immunostaining characterization.* Immunostaining analysis of both bioprinted single cells and spheroids after 7 days of culture showed that no folliculogenesis occurred since both Phalloidin and ZO-1 staining did not underline the presence of any cavities within the cell clusters, which is the main characteristic of thyroid follicles. At the same time, TG expression was low for both single cells and spheroids and no T4 was detected (**Supplementary Figure S4 and Figure 2C and 2D**). These results are consistent with what observed by Kurashige et al. [28] related to the inability of this specific cell line to uptake iodine. Even if this immortalized cell line is important for *in vitro* models, it cannot be fully compared to normal thyrocytes. For this reason, the use of fully differentiated and active thyroid follicles is necessary to produce a physiologically relevant *in vitro* model. Nonetheless, the possibility to bioprint thyroid spheroids shows that our systems could be used in recapitulating and understanding thyroid tumor microenvironment. Thyroid spheroids derived from normal cell lines or cancer cell lines have already been used and clear differences have been shown between 2D and 3D cultured cells in terms of activity and drug response: spheroids presented altered expression of cytoskeletal protein, thyroid differentiation and decreased proliferation [29, 30] compared to cells cultured in classic monolayer.

#### mESCs-derived thyroid follicles bioprinting and characterization

mESCs-derived thyroid follicles were produced as described in **Figure 3A**. When their differentiation was confirmed by morphology and by Tg promoter driven-GFP expression, mESCs-derived follicles were isolated and mixed with AGf5 bioink. After bioprinting, follicles maintained high viability within bioprinted constructs for the first 10 days of culture (**Figure 3B**). We also observed that GFP expression was maintained for the duration of the culture, cells appeared to proliferate over time and assembled into bigger cell clusters. A macroscopic overview of the bioprinted constructs showed that it was still possible for GFP positive follicles to fuse together and to interact with the surrounding cells. Non-thyroid cells (GFP negative cells) proliferated at higher rate than GFP positive thyroid follicles. We also analyzed thyroid marker expression and observed the presence of central lumen, thus proving that both bioinks formulation and bioprinting processing allowed cells to maintain a physiological morphology while inside the bioprinted constructs. Previous studies showed that follicle functionality is strictly correlated to their 3D morphology [21, 31], which is also related to their ability to uptake iodine. Iodine uptake is essential for thyroid hormone generation. Our findings showed that bioprinted follicles expressed both Tg and T4, the first localized mainly in the intracellular compartment, the latter observed inside the lumen created by the follicular cells (**Figure 4**). These results proved that bioprinting had no negative effect on mESCs-derived follicles in terms of viability and functionality. Most importantly, follicles could be maintained for 10 days inside Agf5 bioink, without signs of dedifferentiation and the bioink offered mechanical support to the cells, while allowing cell proliferation and interactions within bioprinted fibres.

**Figure 3.**
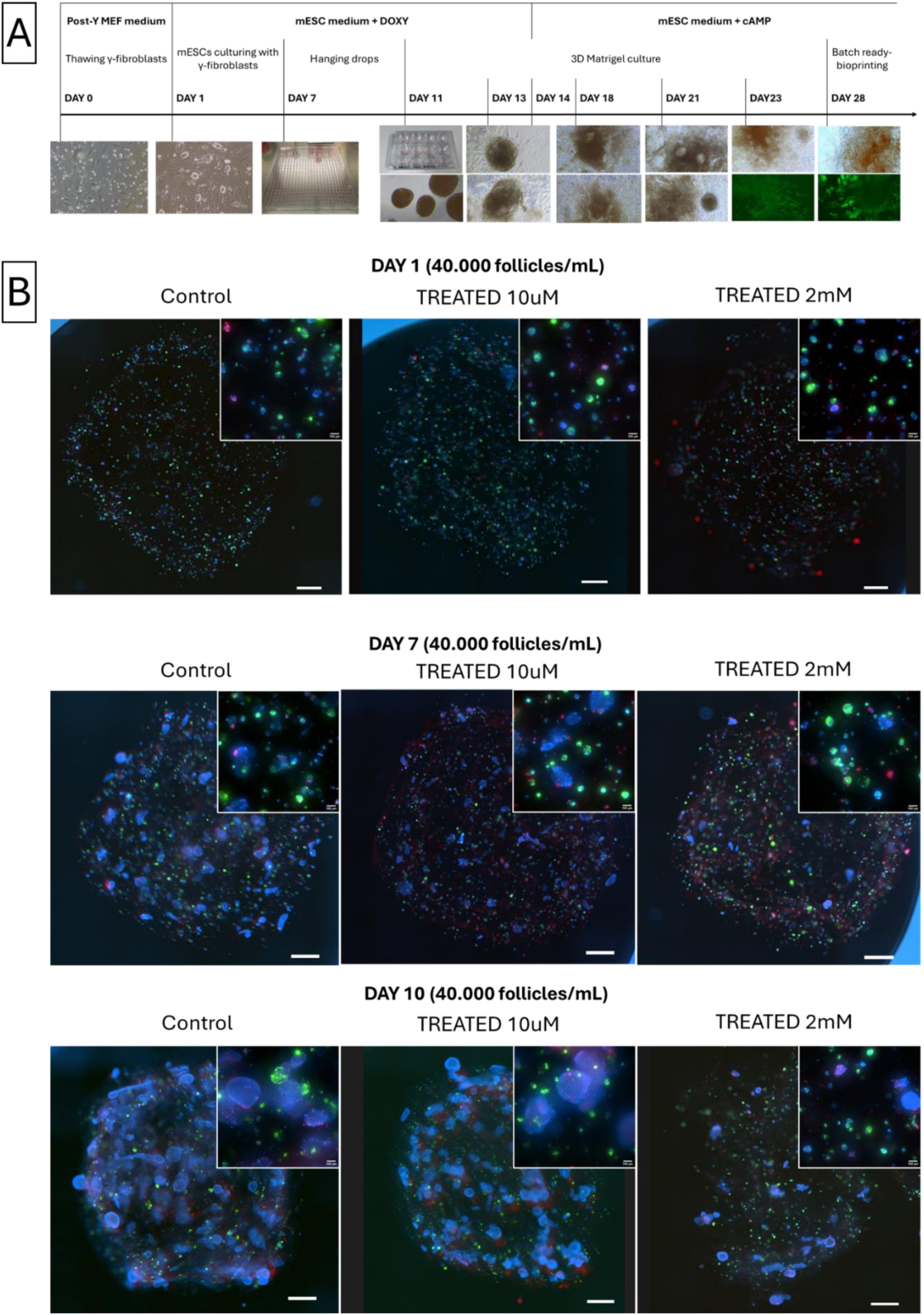
mESC-derived thyroid follicle generation and bioprinting. (**A**) Differentiation overview of mESCs into mESC-derived thyroid follicles inside Matrigel using the protocol developed by Antonica et al[21]. mESCs have been cultured on γ-irradiated fibroblasts for 7 days before being collected and maintained as hanging drops for 5 days. At Day 9 embryonic bodies were collected and embedded in Matrigel for 18 days. Around Day 21 of culture GFP expression was observed showing the commitment to thyroid fate and the co-expression of Nkx2-1 and Pax8. (**B**) Cell distribution and viability of mESC-derived follicles inside bioprinted constructs at day 1, 7 and 10 after bioprinting while exposed to different concentrations of PTU. Since cells were expressing GFP (green), Hoechst was used in order to visualize the nuclei (blue) and Ethidium bromide was used to visualize dead cells (red) (Scale bar:1000 µm-bottom and 100 µm-top).

**Figure 4.**
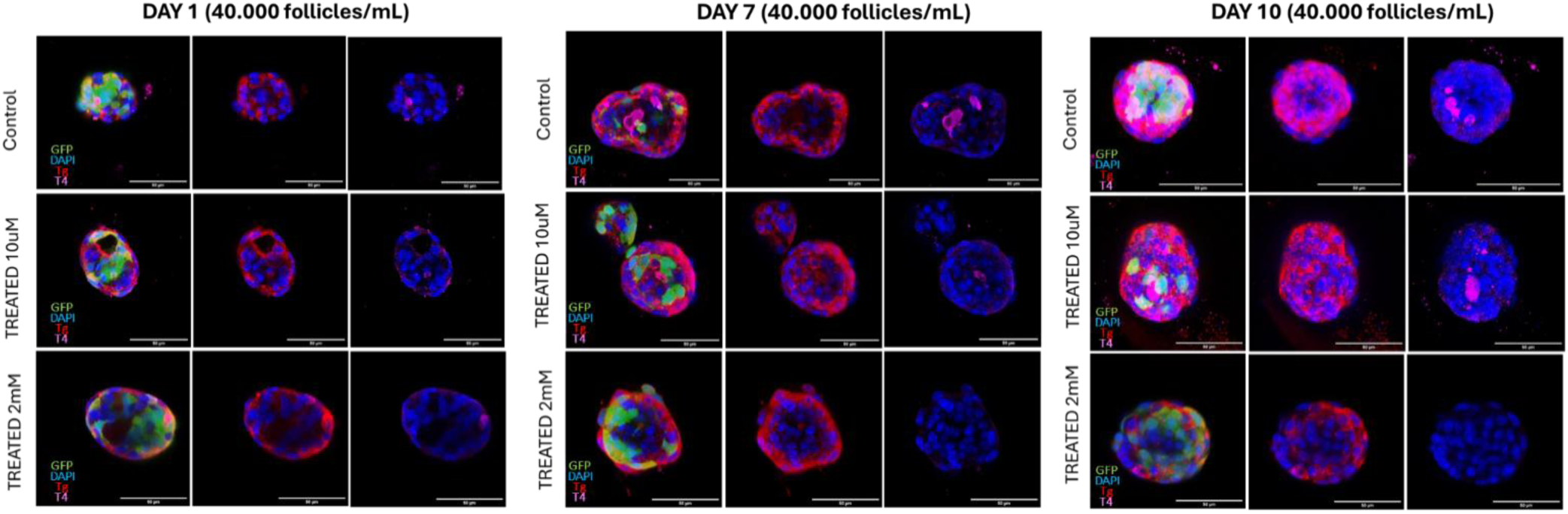
mESCs characterization and PTU effect on T4 release. Bioprinted mESC-derived thyroid follicles immunostaining at day 1, 7 and 10 after bioprinting with different markers: Thyroglobulin (Tg)-red; Thyroxine (T4)-pink; Tight junction protein 1 (ZO-1)-white; DAPI-blue (Scale bar:50µm). **(B)** Boxplot analysis of fold change of T4 release in media for bioprinted constructs treated with two different concentrations of PTU (10 uM=LOW and 2 mM=HIGH) at day 3 and 9 after bioprinting. mESCs-derived thyroid follicles showed no difference in T4 release at day 3 and a significant difference only for 2 mM treatment at day 9 after bioprinting.

### 2.3. mESC-derived follicle exposure to Endocrine disruptor

The ability of the 3D *in vitro* thyroid model to respond to external stimuli was tested by exposing the bioprinted thyroid constructs to PTU, a known ED, at a low (10 µM) and high (2mM) concentration. First, we observed that thyroid differentiation and viability were maintained also after exposure with PTU for a total of 10 days (**Figure 3B**). At different time point after exposure, the expression of GFP and Tg was observed in both untreated and treated follicles, while T4 could be observed only in untreated follicles and follicles treated with the lower concentration of PTU (**Figure 4**). We further assessed if the exposure to PTU could influence the T4 release in the culture media. We observed no difference between the different conditions after 3 days of exposure, while after 9 days only the high concentration of PTU (2 mM) showed a decrease of around 70% in T4 release in the media, thus confirming what observed by T4 immunostaining analysis. **(Figure 5)**.

**Figure 5.**
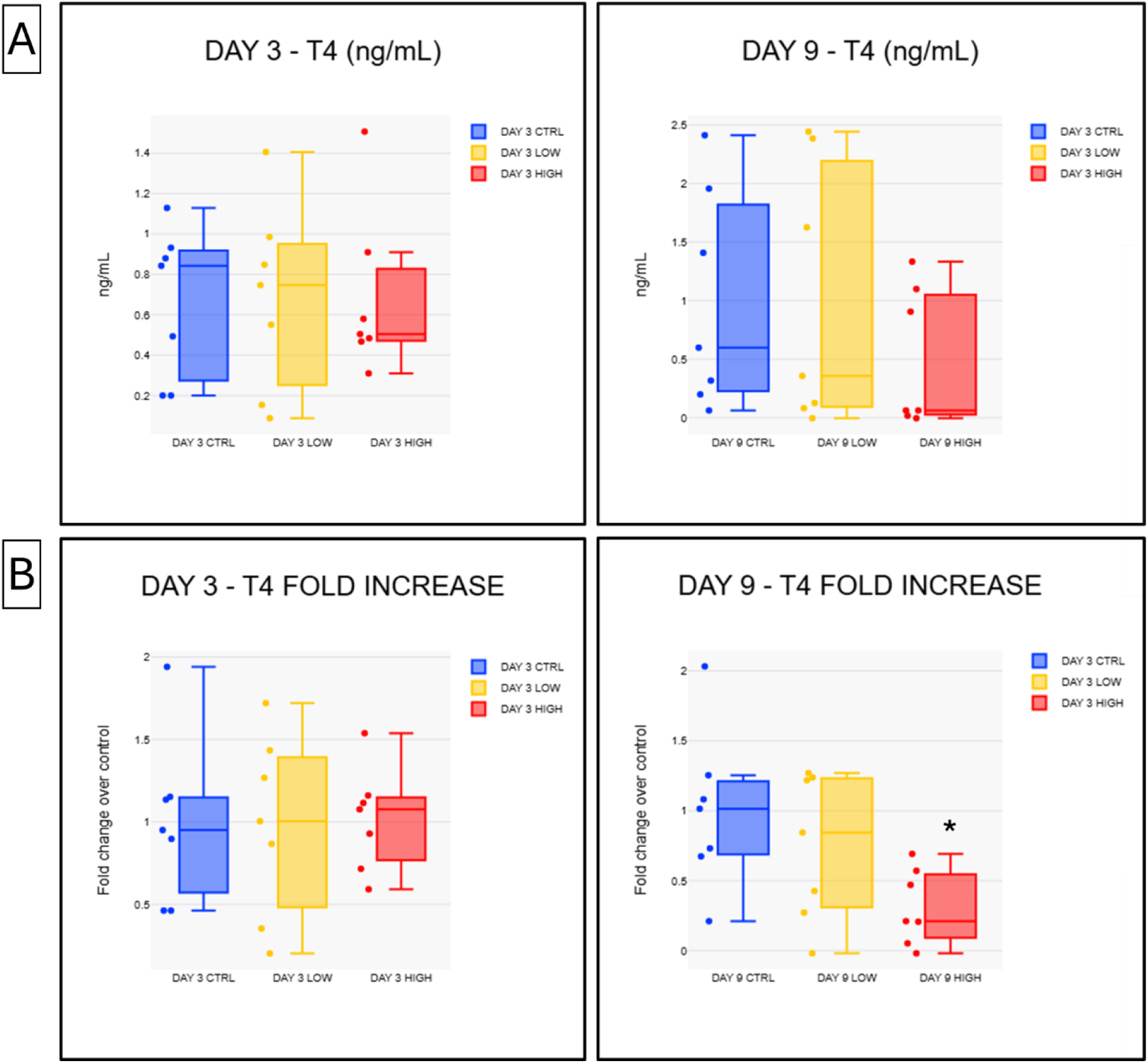
(A)Concentration of T4 released in media and (B) boxplot analysis of fold change of T4 release in media. (N=2, total biological replicates=7) for bioprinted constructs treated with two different concentrations of PTU (10 uM=LOW and 2 mM=HIGH) at day 3 and 9 after bioprinting. mESC-derived thyroid follicles showed no difference in T4 release at day 3 and a significant difference (**\***=p value ≤ 0,05) only for 2 mM treatment at day 9 after bioprinting.

These results show that it was possible to evaluate T4 hormone production by quantifying the level of the hormone released in the media. This is a further prove that our model is not only active, but also able to respond to the exposure to PTU by impairing T4 generation, aligning with PTU effect *in vivo*.

## Discussion

In the last decades, several 3D *in vitro* models of endocrine glands have been developed in order to recapitulate the complexity of gland physiology. The creation of reliable 3D in vitro models can have different purposes ranging from the study of organ development or cancer pathogenesis to the creation of transplantable organs.

In this work, we investigate the creation of a 3D *in vitro* model of the thyroid gland that could be used for the screening of possible EDs and understanding of their effect. Specifically, we evaluated if microfluidic bioprinting could be used to produce a functional thyroid *in vitro* model. Microfluidic bioprinting has already been used for the creation of a kidney [10, 32, 33] and testicular tubules 3D *in vitro* models [34], among other tissues [8, 11]. While the only reported bioprinted thyroid gland construct focused on the production of spheroids obtained from thyroid and allantoic tissue explants [35], our study represents the first use of microfluidic bioprinting for the creation of the first reported thyroid 3D *in vitro* functional model. Bulanova’s work controlled spatial deposition using an extrusion 3D bioprinter using a collagen hydrogel and their ability to fuse over time [35]. Our objective was to harness the specific advantages offered by microfluidic bioprinting, compared to other techniques. Contrary to extrusion bioprinting, microfluidic bioprinting allows the use of low viscosity materials at low concentrations, thus producing soft 3D constructs which will more closely resemble soft gland tissue. Moreover, the combination of low viscosity bioink and low bioprinting pressures ensure a more amenable environment during bioprinting and higher viability. These factors were crucial in choosing microfluidic bioprinting to produce a thyroid *in vitro* model, especially when considering to bioprint delicate cells or complex structures, such as embryonic stem cell-derived thyroid follicles. Finally, this strategy holds the potential to ensure the high-throughput production of 3D constructs containing high concentrations of geometrically organized thyroid organoids.

We initially focused on the screening of several alginate based bioinks that could be at the same time cytocompatible and bioprintable. Nthy-ori 3-1 thyroid immortalized follicular epithelial cells were used to initially test bioinks and the bioprinting procedure. Cells could be maintained inside the bioinks while maintaining viability and metabolic activity, proved by their stable ATP content and their autonomous cluster formations. At the same time, each bioink could be used to produce bioprinted fibres with dimensions ranging from 150 µm to 300 µm using pressure between 20 and 150 mbar, factors that are essential to ensure cell viability during bioprinting and following culture within bioprinted constructs. Nthy-ori 3-1 bioprinted cells had higher viability compared to cells maintained in droplets, while still maintaining stable ATP levels and self-assembly capacities. The increase in cell viability could be caused by the geometry of the bioprinted constructs and the achieved fibre dimension of 200 µm, which allows a more even distribution of the cells and a better diffusion of oxygen and nutrients inside bioprinted fibres compared to the droplets.

This methodology also brings some limitations that need to be considered. The microfluidic bioprinter RX1™ platform was developed to use alginate as a bioink. Alginate is a widely used hydrogel, with rapid crosslinking characteristics and good mechanical properties. Nonetheless, it is not bioactive and requires the presence of other molecules of functionalization to prove an optimal environment for the bioprinted cells. These limitations were overcome by mixing alginate with other molecules. The final bioink composition comprised of alginate and fibrinogen, a protein known for its low immunogenicity, biocompatibility and for favoring cell attachment. A different bioink composed of alginate and fibrinogen has been already used for neural tissues which were able to maintain viability and differentiation after bioprinting [8]. Microfluidic bioprinting is performed in wet conditions while using low viscosity bioinks, factors that can cause the progressive loss of geometric fidelity when bioprinting bigger structures. Nonetheless, the microfluidic bioprinting approach also allowed the high throughput production of reproducible bioprinted constructs with an optimal cell distribution within the constructs, which also maintained their stability for over a week of culture. These characteristics make our microfluidic bioprinting approach an appealing technology to test potential effects of chemical compounds.

Using this bioprinting setup and bioink, not only single cells, but also spheroids produced from Nthy-ori 3-1 cells could be successfully bioprinted. Spheroid bioprinting can suffer from several drawbacks related to the aggregation of spheroids during bioprinting, which could cause clogging inside the printhead or inside the tubing systems [36, 37]. Spheroids and cell aggregates could also lose their morphology and cohesion during bioprinting procedure due to shear forces that might disrupt larger multicellular structures. They could also suffer of higher cell death after bioprinting in case nutrients and oxygen were unable to reach the center of the spheroids, thus causing a necrotic core [38]. None of these problems were detected using our approach and the AGf5 bioink. Spheroids could be bioprinted using different concentrations and were extruded inside bioprinted filaments independently from their diameter. Spheroids not only maintained their cohesiveness after bioprinting, but no necrotic core was observed. Viability remained high up to 7 days after bioprinting.

Despite the optimal results obtained using Nthy-ori 3-1 cells, their characterization after bioprinting showed that both bioprinted spheroids and self-assembled cell clusters obtained from bioprinted single cells did not present follicular morphology, since no lumens were observed and Tg expression was low. Even if this cell line was useful for assessing bioprinting feasibility and bioink cytocompatibility, a more physiologically relevant cell population was necessary for the creation of a functional thyroid 3D *in vitro* model. Despite Nthy-ori 3-1 cell line proved itself to be a useful tool for *in vitro* testing, recent comparison with other thyroid cell lines showed that Nthy-ori 3-1 cell ultimately fail to fully recapitulate thyroid function because of their low or inconsistent ability to uptake iodine or to express thyrotropin receptor (TSHR) and sodium/iodine symporter (NIS)[28]. For this reason, mouse thyroid follicles generated following an already established protocol [21, 22] were used. mESC-derived follicles proved to recapitulate not only follicle morphology, but also functional features, such as expressing Tg and T4 hormone.

We investigated if these characteristics could be maintained after bioprinting and after culture within the novel AGf5 bioink. When assessing the behavior of bioprinted follicles after bioprinting, it was observed that they maintained not only high viability, but also the characteristics of fully differentiated and functional thyroid tissue. Follicles maintained the expression of both Tg and T4 for more than 10 days after bioprinting. Another significant result of this study regards the ability of the 3D *in vitro* thyroid model to react to the exposure to EDs. When exposed to different PTU concentrations, the bioprinted model was able to respond by altering in a significant way the level of T4 hormone released in the cell culture media. These results are consistent with other works that investigated the effect of PTU on rat thyroid cells reporting its effect on T4 levels, thus validating the efficacy and sensitivity of the bioprinted 3D *in vitro* model. It is important to underline that these works observed the effect of PTU *in vivo* [13] or using thyroid follicles obtained from fresh thyroid explants and encapsulated in collagen droplets [39]. To our knowledge, this is the first 3D bioprinted thyroid model able to produce an ED reactive thyroid model from mouse embryonic stem cells, which brings us a step closer to reduce animal testing for ED testing.

## Conclusions

We evaluated the possibility to use microfluidic bioprinting to produce a 3D thyroid model that could have high significance for the investigation of thyroid function and thyroid disruption *in vitro*.

We reported that microfluidic bioprinting can successfully manufacture bioprinted constructs composed of different bioinks combined with immortalized thyroid cells or thyroid follicles obtained from mouse embryonic stem cells. Our findings proved that bioprinted thyroid follicles maintained viability, functionality and differentiation after bioprinting using our homemade bioink. We also proved that microfluidic bioprinting can produce constructs with precise geometries in a high-throughput manner. Bioprinting allowed the production of high density arrays and equidistant deposition of functional thyroid follicles Furthermore, we showed that bioprinted follicles responded to exposure to PTU by reducing the amount of hormone released in the culture media after a chronic exposure of 10 days. Considering the lack of functional 3D thyroid *in vitro* models, these promising results prove that this innovative bioprinted model could be used in the future for the evaluation of potential EDs and their mode of action instead of animal models. Thanks to the controlled spatial alignment of thyroid follicles, such bioprinted thyroid models could be used in the future with other bioprinted organ analogues to study multi-organ response.

## Acknowledgements.

This study was financially supported by the European Union’s Horizon 2020 Research and Innovation Programme under grant agreement no. 825745 (SCREENED).

## Supplementary Information

**Supplementary Table S1.** Biomaterials and features for bioprintability test.

| <u>Biomaterial</u> | <u>Type</u> | <u>Features</u> |
| --- | --- | --- |
| Alginate (Sigma-Aldrich) | Natural-derived | good gelation - good printability - biologically inert - already used in combination with different biomaterials |
| AG10 Matrix (AspectBiosystems) | Natural-derived | modified sodium alginate solution - optimized for AspectBiosystems bioprinter - demonstrated high cell viability and function when bioprinting different cell types |
| Gelatin | Natural-derived | widely used - biocompatible and biodegradable - low antigenicity - can be functionalized - thermoresponsive - gelation is typically slow and unstable (need modification) |
| GelMA | semi-synthetic | functionalized form of Gelatin that enables crosslinking by UV light exposure - provide a suitable cell microenvironment – sufficient integrity but weak mechanical strength |
| Fibrinogen/Fibrin | Natural-derived | biocompatible - biodegradable - non-immunogenic - induce cell attachment, proliferation and ECM formation - used to fabricate microvascular networks - with PEG or a PEG-Gelatin mixture significantly increased degradation time and improved the robustness of the constructs |

**Figure S1.**
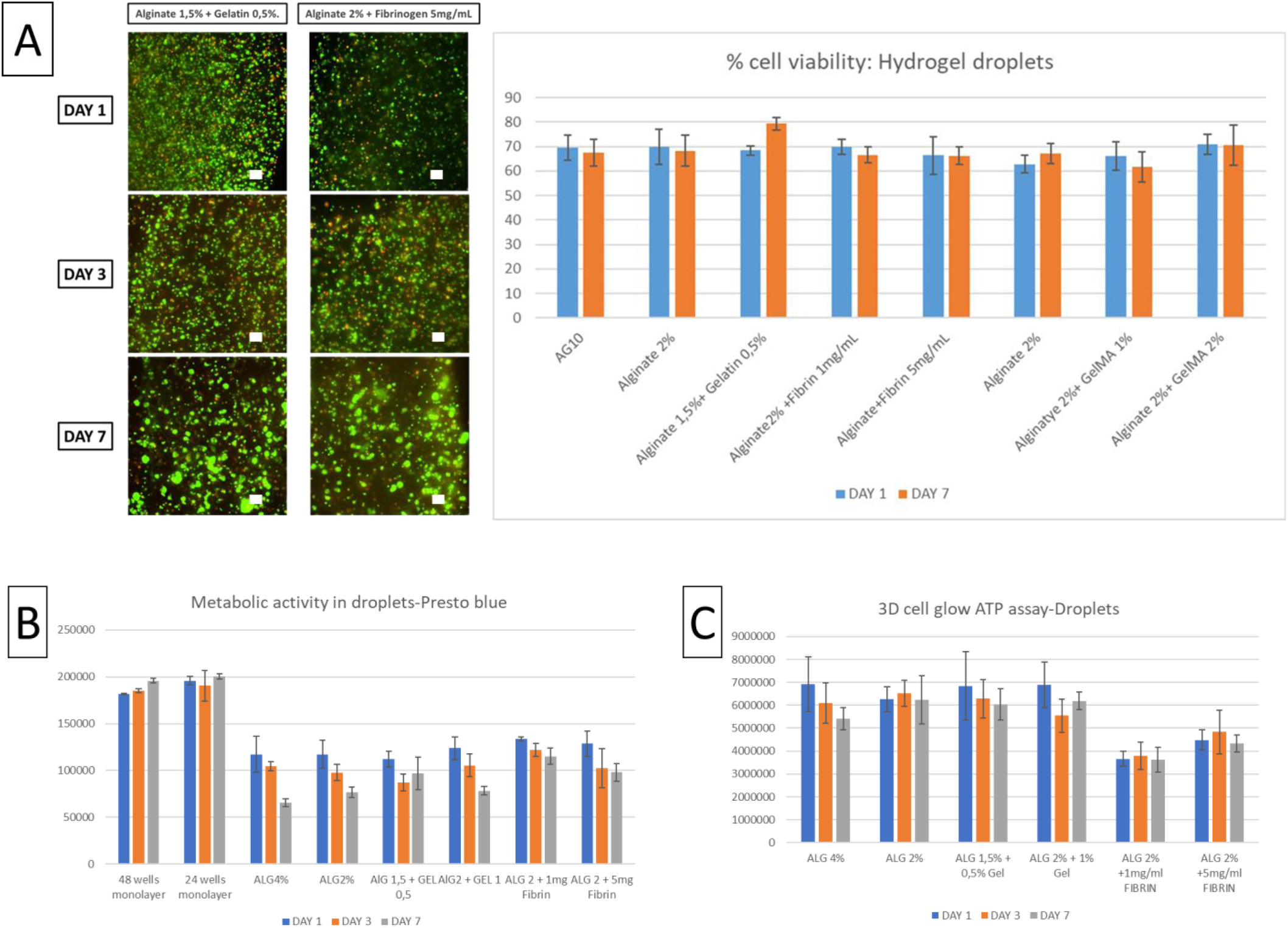
Nthy-ori 3-1 cytocompatibility with different hydrogel compositions. **(A)** Cell viability of Nthy-ori 3-1 encapsulated inside different biomaterial droplets assessed using LIVE/DEAD viability/cytotoxicity Kit and ATP levels assessed using CellTiter-Glo® 3D Assay (Scale Bar: 100µm). **(B)** Metabolic activity assessed by Presto Blue assay from day 1 to day 7 of Nthy-ori 3-1 cultured in 2D monolayers and inside droplets of different hydrogels. **(C)** ATP levels assessed using CellTiter-Glo® 3D Assay from day 1 to day 7 of Nthy-ori 3-1 cultured inside droplets of different hydrogels.

**Figure S2.**
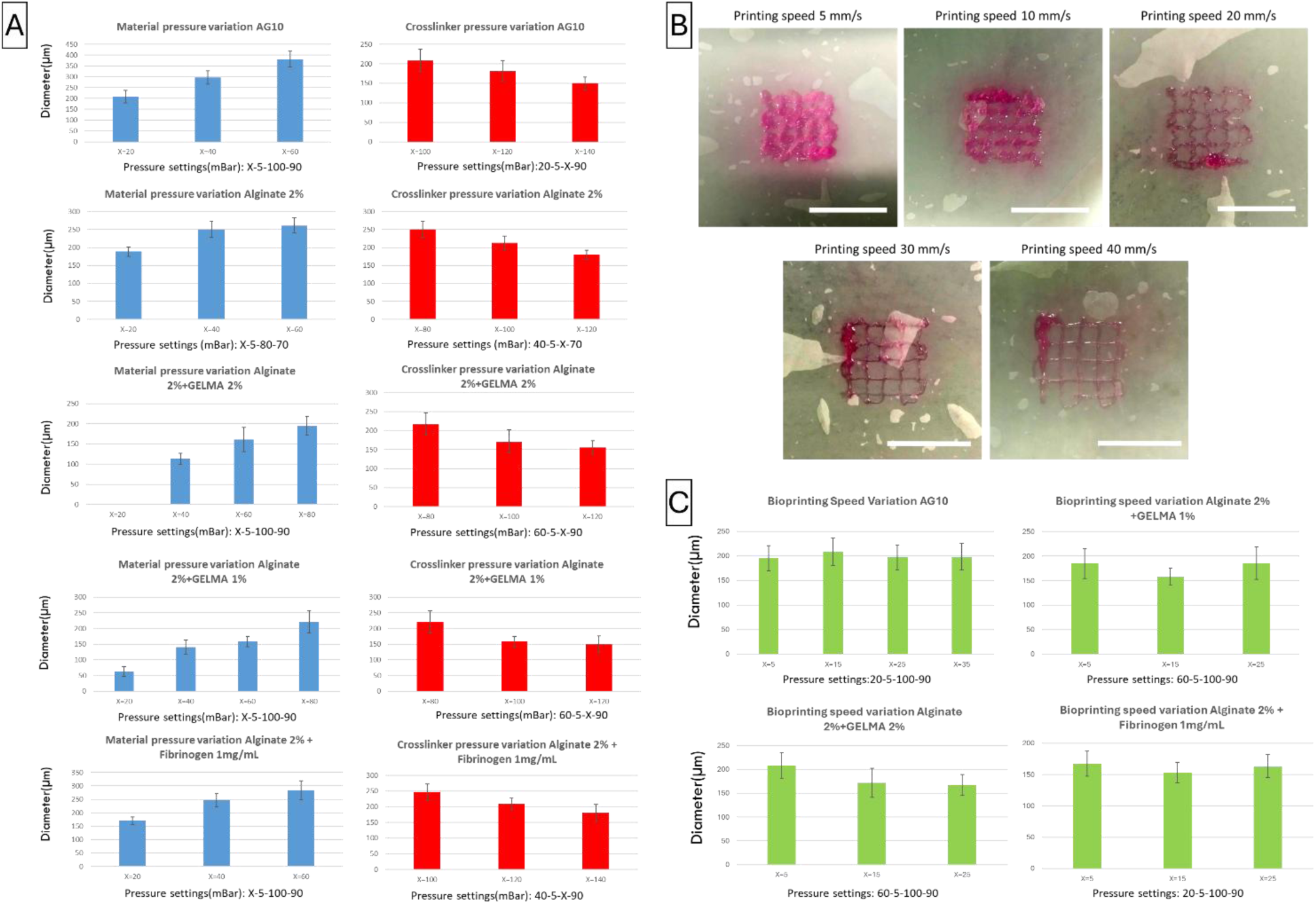
Bioprinting parametric optimization of different bioinks. **(A)** Fibre average diameter obtained by tuning the pressure applied to the material (**BLUE**) or to the crosslinker (**RED**) solution using different bioink compositions. (B) Macroscopic view of bioprinting speed effect on fibre deposition. Scale bar: 10 mm and **(C**) fibre average diameter obtained by tuning bioprinting speed using different bioink compositions.

**Figure S3.**
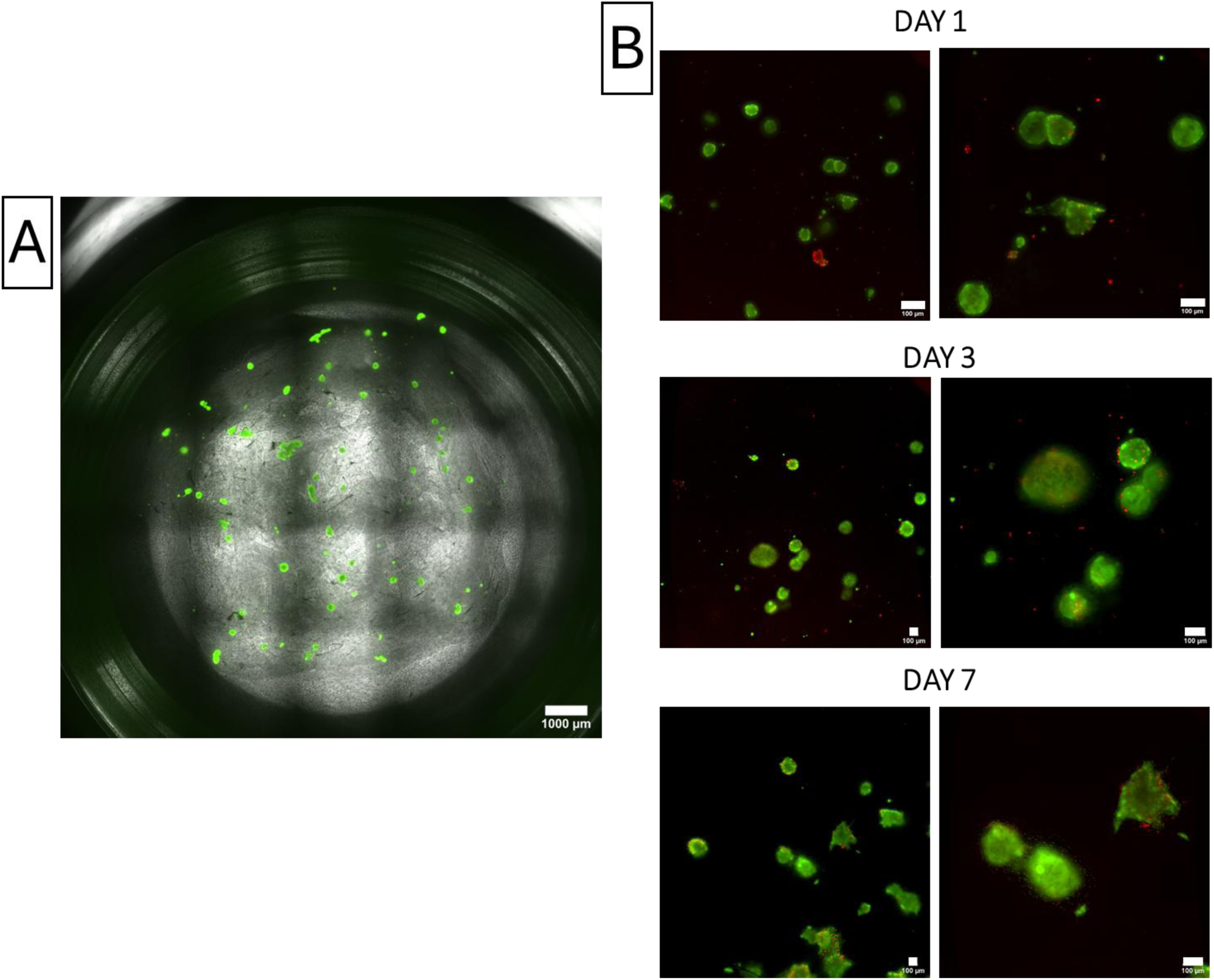
N-thy-ori-3.1 spheroid bioprinting at lower concentration. **(A)** Macroscopic view of spheroid distribution inside bioprinted constructs produced using AGf5 with a final spheroid concentration of 2000 spheroids/mL (Scale bar: 1000 µm). **(B)** Nthy-ori 3-1 spheroids viability assessed on day 1, 3 7 after bioprinting using LIVE/DEAD viability/cytotoxicity kit. Live cells-green, dead cells-red (Scale Bar: 100µm).

**Figure S4.**
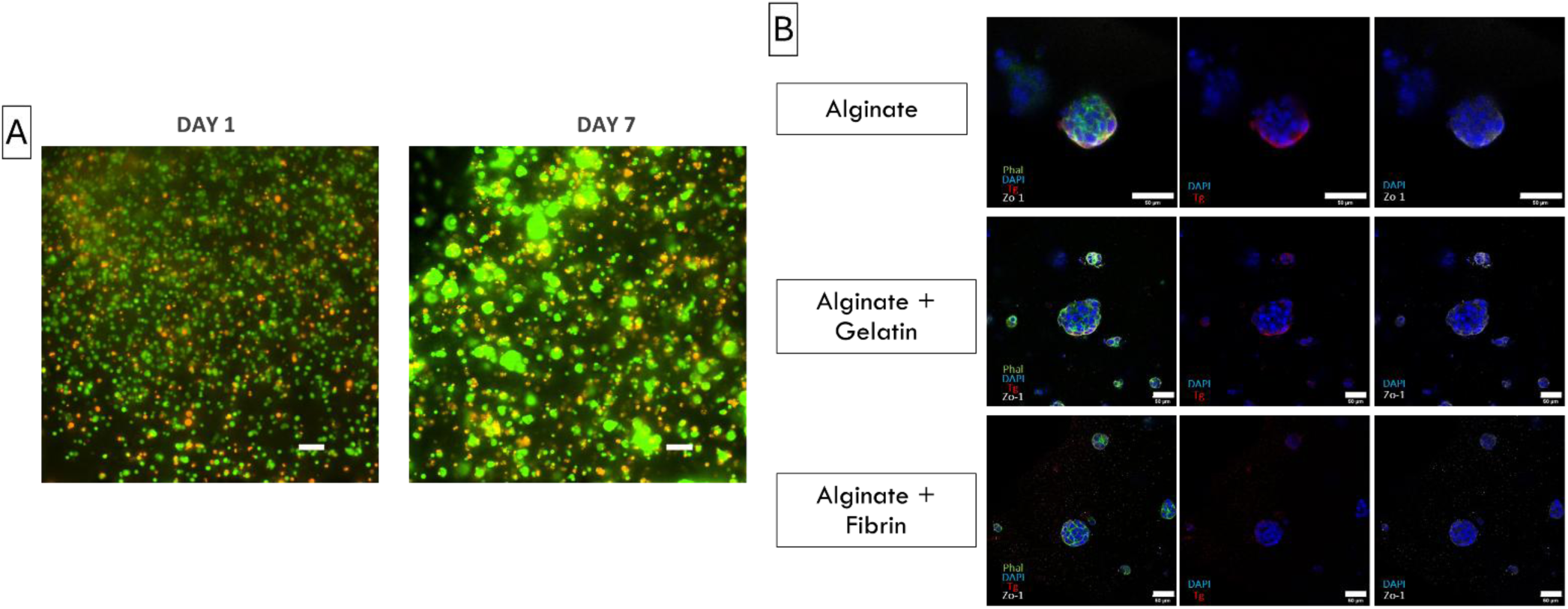
Nthy-ori 3.1 characterization inside hydrogel droplets. **(A)** Cell viability of Nthy-ori 3-1 encapsulated inside AG2 bioink using LIVE/DEAD viability/cytotoxicity Kit (Thermofisher). (B) Nthy-ori 3-1 immunostaining with different markers on day 7 after encapsulation inside AG2, Agg and AGf5 hydrogel droplets: Thyroglobulin (Tg)-red; Thyroxine (T4)-pink; Tight junction protein 1 (ZO-1)-white; Phalloidin-green; DAPI-blue (Scale bar:50 µm).

## References

1. Kirsten, D., The thyroid gland: physiology and pathophysiology. Neonatal Netw, 2000. 19(8): p. 11–26.

2. Stathatos, N., Thyroid physiology. Med Clin North Am, 2012. 96(2): p. 165–73.

3. Stathatos, N., Anatomy and Physiology of the Thyroid Gland: Clinical Correlates to Thyroid Cancer. Thyroid Cancer, September 2016.

4. Moroni, L., et al., SCREENED: A Multistage Model of Thyroid Gland Function for Screening Endocrine-Disrupting Chemicals in a Biologically Sex-Specific Manner. Int J Mol Sci, 2020. 21(10).

5. Slama, R., et al., Scientific Issues Relevant to Setting Regulatory Criteria to Identify Endocrine-Disrupting Substances in the European Union. Environ Health Perspect, 2016. 124(10): p. 1497–1503.

6. OECD, New Scoping Document on in vitro and ex vivo Assays for the Identification of Modulators of Thyroid Hormone Signalling. OECD Publishing, 2017.

7. Colosi, C., et al., Microfluidic Bioprinting of Heterogeneous 3D Tissue Constructs. Methods Mol Biol, 2017. 1612: p. 369–380.

8. Abelseth, E., et al., 3D Printing of Neural Tissues Derived from Human Induced Pluripotent Stem Cells Using a Fibrin-Based Bioink. ACS Biomater Sci Eng, 2019. 5(1): p. 234–243.

9. Robinson, M., et al., Microfluidic bioprinting for the in vitro generation of novel biomimetic human testicular tissues. 2021.

10. Addario, G., et al., Microfluidic bioprinting towards a renal in vitro model. Bioprinting, 2020. 20.

11. Davoodi, E., et al., Extrusion and Microfluidic-based Bioprinting to Fabricate Biomimetic Tissues and Organs. Adv Mater Technol, 2020. 5(8).

12. Colosi, C., et al., Microfluidic Bioprinting of Heterogeneous 3D Tissue Constructs Using Low-Viscosity Bioink. Adv Mater, 2016. 28(4): p. 677–84.

13. Hassan, I., et al., Extrapolating In Vitro Screening Assay Data for Thyroperoxidase Inhibition to Predict Serum Thyroid Hormones in the Rat. Toxicol Sci, 2020. 173(2): p. 280–292.

14. Lemoine, N.R., et al., Characterisation of human thyroid epithelial cells immortalised in vitro by simian virus 40 DNA transfection. Br J Cancer, 1989. 60(6): p. 897–903.

15. Li, X., et al., Proteomic analysis of differentially expressed proteins in normal human thyroid cells transfected with PPFP. Endocr Relat Cancer, 2012. 19(5): p. 681–94.

16. Rajoria, S., et al., Metastatic phenotype is regulated by estrogen in thyroid cells. Thyroid, 2010. 20(1): p. 33–41.

17. Wynford-Thomas, D., et al., Conditional immortalization of human thyroid epithelial cells: a tool for analysis of oncogene action. Mol Cell Biol, 1990. 10(10): p. 5365–77.

18. Kim, B.A., et al., Expression Profiling of a Human Thyroid Cell Line Stably Expressing the BRAFV600E Mutation. Cancer Genomics Proteomics, 2017. 14(1): p. 53–67.

19. Carvalho, D.J., et al., Thyroid-on-a-Chip: An Organoid Platform for In Vitro Assessment of Endocrine Disruption. Adv Healthc Mater, 2023. 12(8): p. e2201555.

20. Fois, M.G., et al., Assessment of Cell-Material Interactions in Three Dimensions through Dispersed Coaggregation of Microsized Biomaterials into Tissue Spheroids. Small, 2022. 18(29): p. e2202112.

21. Antonica, F., et al., Generation of functional thyroid from embryonic stem cells. Nature, 2012. 491(7422): p. 66–71.

22. Romitti, M., et al., Single-Cell Trajectory Inference Guided Enhancement of Thyroid Maturation In Vitro Using TGF-Beta Inhibition. Front Endocrinol (Lausanne), 2021. 12: p. 657195.

23. Yoshihara, A., et al., Inhibitory effects of methimazole and propylthiouracil on iodotyrosine deiodinase 1 in thyrocytes. Endocr J, 2019. 66(4): p. 349–357.

24. Cui, H., et al., 3D Bioprinting for Organ Regeneration. Adv Healthc Mater, 2017. 6(1).

25. Carmeliet, P. and R.K. Jain, Angiogenesis in cancer and other diseases. Nature, 2000. 407(6801): p. 249–57.

26. Lovett, M., et al., Vascularization strategies for tissue engineering. Tissue Eng Part B Rev, 2009. 15(3): p. 353–70.

27. Ozbolat, I.T. and M. Hospodiuk, Current advances and future perspectives in extrusion-based bioprinting. Biomaterials, 2016. 76: p. 321–43.

28. Kurashige, T., M. Shimamura, and Y. Nagayama, Reevaluation of the Effect of Iodine on Thyroid Cell Survival and Function Using PCCL3 and Nthy-ori 3-1 Cells. J Endocr Soc, 2020. 4(11): p. bvaa146.

29. Oh, J.M., et al., Different Expression of Thyroid-Specific Proteins in Thyroid Cancer Cells between 2-Dimensional (2D) and 3-Dimensional (3D) Culture Environment. Cells, 2022. 11(22).

30. Lee, M.A., et al., Novel three-dimensional cultures provide insights into thyroid cancer behavior. Endocrine-Related Cancer, 2020. 27(2): p. 111–121.

31. Toda, S., et al., Culture models for studying thyroid biology and disorders. ISRN Endocrinol, 2011. 2011: p. 275782.

32. Perin, F., et al., Bioprinting of Alginate-Norbornene bioinks to create a versatile platform for kidney in vitro modeling. Bioact Mater, 2025. 49: p. 550–563.

33. Fransen, M.F.J., et al., Bioprinting of kidney in vitro models: cells, biomaterials, and manufacturing techniques. Essays Biochem, 2021. 65(3): p. 587–602.

34. Robinson, M., et al., Microfluidic bioprinting for the in vitro generation of novel biomimetic human testicular tissue. 2021.

35. Bulanova, E.A., et al., Bioprinting of a functional vascularized mouse thyroid gland construct. Biofabrication, 2017. 9(3): p. 034105.

36. Ng, W.L., et al., Microvalve-based bioprinting - process, bio-inks and applications. Biomater Sci, 2017. 5(4): p. 632–647.

37. Ren, Y., et al., Developments and Opportunities for 3D Bioprinted Organoids. Int J Bioprint, 2021. 7(3): p. 364.

38. Decarli, M.C., et al., Cell spheroids as a versatile research platform: formation mechanisms, high throughput production, characterization and applications. Biofabrication, 2021. 13(3).

39. Clavel Rolland, N., et al., Investigating the mechanisms of action of thyroid disruptors: A multimodal approach that integrates in vitro and metabolomic analysis. Toxicol In Vitro, 2024. 100: p. 105911.

